# Phase-Specific Hippocampal and Cortical Medial Temporal Lobe Involvement in Allocentric Working Memory

**DOI:** 10.1101/2025.11.06.686935

**Authors:** Emilie A. Orvik, Anders M. Fjell, Knut Øverbye, Kristine B. Walhovd, Markus H. Sneve, Håkon Grydeland

**Affiliations:** Center for Lifespan Changes in Brain and Cognition, Department of Psychology, University of Oslo, Forskningsveien 3A, 0373 Oslo, Norway; Computational Radiology and Artificial Intelligence, Department of Radiology and Nuclear Medicine, Oslo University Hospital, Sognsvannsveien 20, 0424 Oslo, Norway

**Keywords:** allocentric working memory, hippocampus, medial temporal lobe, fMRI, aging, relational binding

## Abstract

The hippocampus and cortices of the medial temporal lobe (MTL) are increasingly implicated in working memory, particularly for tasks requiring complex, high-resolution relational binding, but their precise contributions—particularly during delay maintenance—remain debated. Using functional magnetic resonance imaging (fMRI), we investigated the involvement of hippocampal and cortical MTL in a demanding allocentric working memory task requiring high-resolution, relational binding. During fMRI, 128 healthy human adults (92 females) aged 20–83 (mean 39) years performed a task in which object–location bindings had to be learned, maintained, and manipulated across an 8 second delay. The design included a passive viewing condition to control for perceptual and attentional demands and a staircase procedure to balance task difficulty across participants. Across the full sample, anterior and mid hippocampal subregions and cortical MTL areas were engaged during encoding. In the delay phase, we observed a non-uniform pattern, with hippocampal and entorhinal deactivation alongside perirhinal and parahippocampal activation. During test, activation occurred in posterior hippocampus and several cortical MTL regions. Contrary to predictions, hippocampal and cortical MTL activation did not vary with performance among younger adults. Instead, differences emerged in a left temporoparietal cluster, potentially reflecting verbal encoding strategies. Older adults, relative to younger adults with the most comparable performance levels, showed lower anterior hippocampal activation and smaller correct–passive viewing differences in perirhinal and parahippocampal cortices. Taken together, these findings demonstrate distinct phase-specific hippocampal and cortical MTL involvement across the temporal unfolding of allocentric working memory, with evidence of hippocampal engagement even during the delay, yet the absence of MTL-performance associations among younger adults suggests this involvement, while robust, may play a more auxiliary role.

## 1. Introduction

The hippocampus, in addition to its well-established role in long-term memory, has been implicated in working memory processes, that is, the temporary retention and manipulation of information over short periods (Ranganath & D’Esposito, 2001). However, the exact nature of hippocampal involvement remains unclear, particularly during the delay period, when information must be actively maintained in the absence of external input (Sreenivasan & D’Esposito, 2019). While findings from intracranial recordings have demonstrated persistent MTL neuronal activity during working memory delays for both individual visual features and categorical stimuli (Kamiński et al., 2017; Xie et al., 2023), functional magnetic resonance imaging (fMRI) studies in healthy participants have yielded a more ambiguous picture. Specifically, although involvement of both anterior and posterior hippocampus has been reported during working memory (Grady, 2020), uncertainty persists as to whether this reflects engagement also during the delay period – as opposed to transient responses during encoding or retrieval – with some arguing that these hippocampal signals instead reflect long-term memory processes (Slotnick, 2022).

A reconciling proposition is that hippocampus might not be uniformly involved in all kinds of working memory, instead being more strongly involved for tasks that require certain forms or levels of processing. One suggestion poses that hippocampal involvement increases when working memory capacity demands exceed what can be maintained by other regions (Jeneson & Squire, 2012). A related but distinct account emphasizes representational complexity rather than capacity per se. Patient studies have demonstrated that hippocampal damage impairs performance on tasks requiring (i) conjunctions of object identity and spatial location irrespective of retention time frame (Olson et al., 2006), and (ii) visual search for a target among complex stimuli without delays (Warren et al., 2011). More broadly, Yonelinas (2013) argues that the hippocampus plays a critical role specifically in relations or bindings of a complex and high-resolution nature. Crucially, when it comes to the delay period, operational complexity interacts with retention interval in predicting hippocampal criticality: hippocampus uniquely supports complex high-resolution bindings over long delay periods, while, as the delay decreases, such bindings can be supported by cortical regions (Yonelinas, 2013).

Tasks requiring high-resolution relational binding are thus well suited to test hippocampal and cortical MTL involvement in working memory across task phases. Allocentric spatial tasks that require encoding and maintaining precise configurations among multiple objects (Burgess, 2008; Hartley et al., 2004) represent a strong example of such tasks, as they combine viewpoint-independent spatial processing with the kind of complex, high-resolution bindings that Yonelinas (2013) argues are critical for hippocampal involvement. Using such an allocentric task, Hannula and Ranganath (2008) showed that anterior and posterior hippocampus, as well as cortical MTL regions, were activated during successful encoding and retrieval of relational information, but hippocampal involvement was not observed during the delay period. Instead, entorhinal cortex activation in the early phase of the delay correlated with performance accuracy. This pattern was consistent with the fMRI study of Schon et al. (2016), using a complex scene working memory task, which also reported only cortical MTL delay-period activity.

While these studies demonstrate hippocampal and cortical MTL involvement during encoding and retrieval of high-resolution relational information, whether hippocampal engagement extends to the delay period remains to be fully characterized, particularly in larger samples with task designs that control for perceptual demands and accommodate individual differences in ability. Moreover, given the well-documented sensitivity of hippocampal and MTL structures to age-related decline (Nyberg & Pudas, 2019) and the age-related deterioration of allocentric spatial processing (Van Der Ham & Claessen, 2020), examining how these activation patterns differ across the adult lifespan could provide novel clues to the role of hippocampal and cortical MTL in delay maintenance.

Here, we address these questions using a demanding allocentric working memory task requiring high-resolution relational binding of object-location information across an 8-second delay. We tested 128 healthy adults aged 20–83 years, allowing us to examine not only the phase-specific involvement of hippocampal and cortical MTL regions across encoding, maintenance, and retrieval, but also how this involvement relates to individual differences in performance and to aging. To isolate allocentric processing from perceptual and attentional demands, we included a passive viewing condition matched in visual input, scene rotation and response requirements. This design feature allows identification of modulations specific to relational binding beyond shared sensory and motor processing. To enable meaningful comparisons across individuals of differing ability, task difficulty was individually adjusted using a staircase procedure. We hypothesized that hippocampal and cortical MTL regions would be engaged across all three phases, with potentially distinct contributions, and that older adults would show lower engagement particularly in hippocampal regions.

## 2. Methods

### 2.1 Participants

The project was approved by the Norwegian Regional Committee for Medical and Health Research Ethics, South-East. All participants signed informed consent and were compensated for their participation.

Task data were collected from 151 participants, aged 20 to 89 years, drawn from two different samples. The primary sample was recruited through Facebook ads and screened at the Center for Lifespan Changes in Brain and Cognition (Department of Psychology, University of Oslo). To increase the number of older participants, we included a subsample of cognitively healthy individuals who had been recruited as a control group in a study of cognitive decline in delirium patients, with recruitment and screening procedures previously described (Idland et al., 2017). Sample size was determined by participant availability across the two recruitment streams and the practical constraints of MRI data collection (scanner time and project resources). We aimed to include as many eligible participants as feasible within the study period. Participants had no history of neurological or severe psychiatric conditions, major head trauma, psychiatric treatment, or medication known to alter nervous system function. Participants were required to have an intelligence quotient equal to or above 85, as measured with the Wechsler Abbreviated Scale of Intelligence (Wechsler, 1999). Older participants (60+) were required to score ≥ 26 on the Mini-Mental Status Exam (MMSE; Folstein et al., 1975). Additionally, to ensure task comprehension, inclusion required performance above chance level (>0.33) on the fMRI task. A total of 23 participants were excluded for the following reasons: 1 participant above the age of 60 scored below 26 on the MMSE, 13 participants scored below chance level (<0.33) on the task administered in the scanner, 2 participants had missing behavioral data, 1 participant had functional images of insufficient quality, and 6 participants were excluded for excessive head movement, defined as mean relative root-mean-square (RMS) displacement (average frame-to-frame head motion in millimeters, calculated from the six rigid-body realignment parameters with rotations converted to millimeters assuming a 50-mm head radius) exceeding 1.5 times the interquartile range above the upper quartile, calculated separately for each participant group (see Supplemental Figure S1). The final sample consisted of 128 participants (see **Table 1** for participant characteristics), with more women especially among the younger participants (W = 1237, p = .027), and accordingly women’s age distribution was younger than men’s (median 30.8 [IQR 15.6] vs 36.1 [IQR 43.1] years; **Fig. 2A**).

**Table 1.** Participants characteristics. **Abbreviations:** n = sample size ^a^ One younger adult (under age 40) scored below the MMSE cutoff of 26; this criterion only applied to participants over 60. ^b^ IQ scores were not available for participants in Sample 2

|  | Full Sample |  |  | Sample 1 |  |  | Sample 2 |  |  |
| --- | --- | --- | --- | --- | --- | --- | --- | --- | --- |
|  | Mean (SD) | n | Range | Mean (SD) | n | Range | Mean (SD) | n | Range |
| Age | 39.4 (17.8) | 128 | 20.5 - 83.0 | 33.6 (11.0) | 110 | 20.5 - 70.3 | 75.0 (3.9) | 18 | 69.5 - 83.0 |
| Sex (female/male) | - | 92/36 | - | - | 85/25 | - | - | 7/11 | - |
| MMSE | 29.1 (1.1) | 120 | 24.0 - 30.0<sup>a</sup> | 29.1 (1.1) | 104 | 24.0 - 30.0<sup>a</sup> | 29.4 (1.0) | 16 | 27.0 - 30.0 |
| IQ | 115.8 (8.4) | 104 | 96.0 - 132.0 | 115.8 (8.4) | 104 | 96.0 - 132.0 | -<sup>b</sup> | 0<sup>b</sup> | -<sup>b</sup> |
| Education | 16.0 (2.4) | 118 | 8.0 - 24.0 | 16.1 (2.1) | 102 | 12.0 - 22.0 | 15.8 (3.9) | 16 | 8.0 - 24.0 |

Because the middle-aged sample was small and we sought age contrasts at comparable performance, we performed group analyses by defining two age groups, younger (20–40 years, n = 89) and older (60+ years, n = 23), thereby excluding the sixteen middle-aged participants (40 < age < 60) from the group analyses. As shown in **Fig. 2B**, task performance varied substantially among younger individuals. To examine performance-related differences within this group, younger participants were further divided based on their average load level, which reflects the mean number of items they were able to track across trials. A median split (load level = 3.66) was used as the cutoff, creating two subgroups: Lower-performing younger adults (average load level at or below the median, n = 45) and higher-performing younger adults (above the median, n = 44). Because the task dynamically adjusted load to keep difficulty comparable across participants, we defined performance as the average load level achieved (details below). By design, average load level strongly correlated with accuracy (ρ = 0.93, p < 0.001; **Fig. 2C**). There was no significant difference in IQ between the two younger groups (t(82) = -1.17, p = .24). To test for age-related differences, we compared activity between older adults (n = 23, 61-83 years) and younger adults with the most similar performance level, specifically the lower-performing younger adults.

### 2.2 fMRI Task

Participants completed a spatial working memory task (**Fig. 1**), building on the task by Hannula and Ranganath (2008), comprising distinct scenes rendered in Unreal Engine 4 and presented using the E-Prime software. Initially, participants were presented with a study scene for a duration of 8.25 seconds. Subsequently, the objects within the scene were concealed by an ascending brick wall, and the entire scene then rotated 90 degrees in either a clockwise or counterclockwise direction, for an equivalent duration of 8.25 seconds. Upon the descent of the wall and the subsequent reappearance of the objects, participants were to provide one of three potential responses based on the observed configuration: (1) ‘match,’ where the positions of the objects relative to each other remained unaltered, (2) ‘mismatch-position,’ in which one object had been relocated to a different location, or (3) ‘mismatch-swap,’ where two objects had exchanged positions. They had 6.05 seconds to evaluate the test display and respond, followed by a jittered intertrial interval ranging from 3.65 to 11.65 seconds (mean = 6.52 seconds). The task began with scenes containing two objects. After two consecutive correct responses, an additional object was added, up to a maximum of five. Conversely, an incorrect response reduced the number of objects by one. The staircase was designed to converge on 70.7% accuracy under ideal conditions. However, as reported in the Results, some participants performed well above or below this target, indicating that the maximum and minimum difficulty levels did not perfectly match all participants. Control, passive viewing trials featured a single object, namely a spherical rock, which was concealed behind the brick wall and underwent a 90-degree rotation. The object’s position remained unchanged, and the correct response to the control trials would therefore always be ‘match’.

**Figure 1.**
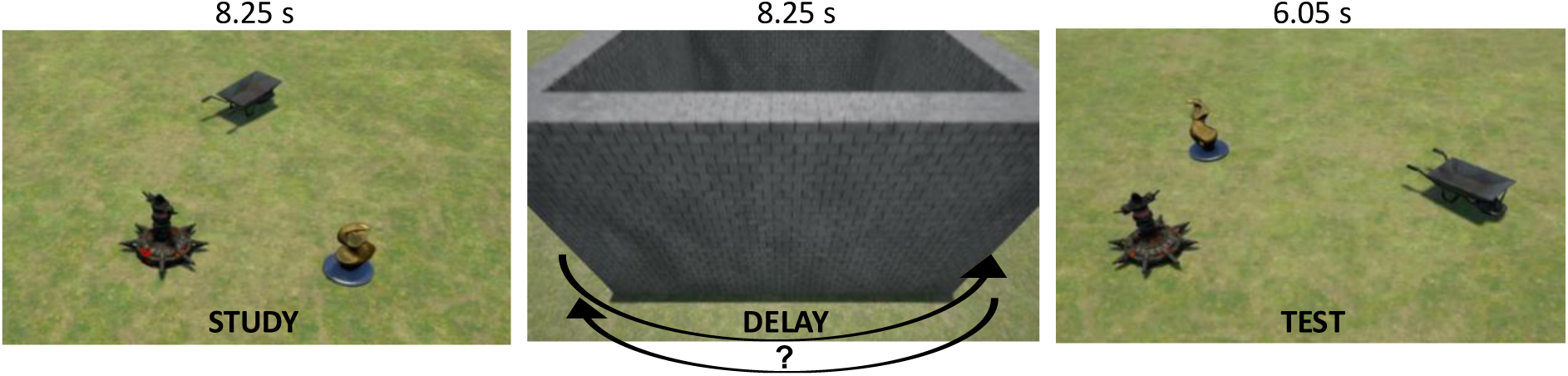
The spatial working memory task, illustrating its three phases. This example depicts a mismatch-swap trial. After the whole scene turns clockwise, the yellow and the black objects have swapped positions. The arrows indicate how the brick wall can turn either clockwise or counterclockwise.

### 2.3 Procedure

Before scanning, participants read the task instructions and viewed three example scenes with a transparent wall, allowing them to observe object movements in the mismatch-position and mismatch-swap conditions, as well as object stability in the match condition. The participants were instructed that each trial would start with them seeing a set of objects. Their job was to always remember the exact position of each object, and after the rotation they had to decide on three possible answers: 1) No change (none of the objects changed position), 2) One object moved to a new position, 3) Two objects swapped places. Notably, they were informed that all three answers were equally likely. Furthermore, participants were instructed that they would sometimes see only a single static object, the spherical rock. They were instructed to remain attentive during these trials, even though the correct response was always “No change”. Lastly, they were informed that the number of objects could vary from trial to trial. After these instructions, participants completed a 10-minute training session outside the scanner, consisting of trials with two objects in the first half and three objects in the second half, as well as passive viewing trials. They were then positioned in the scanner and performed the first run consisting of 15 working memory trials and 5 passive viewing trials. After a short break without exiting or repositioning in the scanner, the second run started with the same number of trials. In this second run, the participants started with the number of objects corresponding to the difficulty level they spent the most time on during the first run, reflecting the level at which they performed most consistently.

### 2.4 MRI Acquisition

Imaging data were collected on a 3T Siemens Prisma MRI system equipped with a 32-channel Siemens head coil (Siemens Medical Solutions, Erlangen, Germany) at Oslo University Hospital Rikshospitalet. Functional images were acquired using a gradient-echo echo-planar imaging (EPI) sequence with a multiband acceleration factor of 8, and with a pixel bandwidth of 2480 Hz. Each functional volume consisted of 56 transversally oriented slices (no gap) with a repetition time (TR) of 550 ms, an echo time (TE) of 33 ms, and a flip angle of 50°. The acquisition matrix was 96 × 96, with an isotropic voxel size of 2.5 mm and a field of view (FOV) of 240 × 240 × 140 mm. The phase encoding direction was anterior-to-posterior. The total readout time was 0.052 s. High-resolution anatomical images were acquired using a 3D Magnetization-Prepared Rapid Gradient-Echo (MPRAGE) sequence for T1-weighted imaging and a 3D SPACE sequence for T2-weighted imaging. Both sequences had an acquisition matrix of 208 × 300 × 320, an isotropic voxel size of 0.8 mm, and a field of view (FOV) of 166.4 × 240 × 256 mm. T1-weighted images were acquired with a repetition time (TR) of 2400 ms, an echo time (TE) of 2.22 ms, and a flip angle of 8°, while T2-weighted images had a TR of 3200 ms, a TE of 563 ms, and a flip angle of 120°. The phase encoding direction was right-to-left for both sequences. The pixel bandwidth was 220 Hz for T1-weighted images and 745 Hz for T2-weighted images. Data were acquired with the participant positioned head-first supine (HFS) using Siemens’ syngo MR E11 software.

### 2.5 Preprocessing

Preprocessing of anatomical and functional MRI data was performed using the Human Connectome Project (HCP) minimal preprocessing pipeline version 4.3.0 (Glasser et al., 2013). These pipelines consist of standardized structural and functional workflows optimized for high-resolution, multiband imaging data.

Anatomical preprocessing followed the HCP structural pipeline, which includes the PreFreeSurfer, FreeSurfer, and PostFreeSurfer stages – responsible respectively for correcting and aligning structural images, performing brain segmentation and cortical surface reconstruction, refining, and converting the results into a format ready for surface-based group analyses. T1-weighted and T2-weighted images were corrected for gradient distortions and intensity inhomogeneities, aligned to the anterior commissure-posterior commissure (AC-PC) axis, and skull stripped. Cortical and subcortical segmentations, as well as cortical surface reconstruction, were generated using FreeSurfer (v6.0; Fischl, 2012). PostFreeSurfer processing refined surface reconstructions using multimodal structural data, generated myelin maps and cortical thickness estimates, and resampled individual anatomies into a standardized surface space (fs_LR 32k mesh) for compatibility with downstream analyses.

Functional MRI preprocessing used the HCP volume and surface pipelines, which included correction for gradient distortions, realignment for within-run head motion, and correction of susceptibility-induced distortions using spin-echo images with opposing phase-encoding directions to estimate field maps (Andersson et al., 2003; Smith et al., 2004). Functional volumes were registered to each participant’s structural image and subsequently normalized to MNI152 standard space. Intensity normalization and bias field correction were also applied. Slice timing correction was not applied, in accordance with the HCP pipeline recommendations. This decision reflects the high temporal resolution enabled by the multiband EPI sequence during acquisition (TR = 550 ms, multiband factor = 8). Functional data were then projected onto the individual cortical surface reconstruction and resampled into fs_LR space. Minimal surface-based smoothing (2mm FWHM) was applied during this step, consistent with HCP pipeline defaults. The final output was stored in the CIFTI grayordinates format, integrating surface-based cortical data and volumetric subcortical data.

After preprocessing, denoising was performed using the ICA-FIX pipeline (Griffanti et al., 2014; Salimi-Khorshidi et al., 2014). Independent component analysis (ICA) was used to identify components reflecting noise sources (e.g., motion, physiological artifacts, scanner-related fluctuations), which were then classified and removed. Denoising was performed using ICA-FIX on both single runs and multi-run concatenated data, following the HCP-style pipeline. To further mitigate artifacts from systemic low frequency oscillations, RapidTide (Frederick et al., 2012; Korponay et al., 2024) was used to estimate and regress out voxelwise delay-adjusted signals reflecting global physiological fluctuations. Delay models were generated using pre-denoising data and applied post-denoising to account for temporally shifted physiological noise sources, such as respiration and cardiac-related fluctuations (Wanger et al., 2023).

### 2.6 Deconvolution and ROI Extraction

fMRI data were analyzed using an event-related deconvolution approach implemented in nideconv (Hollander et al., 2019). The GroupResponseFitter function was used to estimate the hemodynamic response for each event type (correct trials, incorrect trials, and passive viewing), modeling activity over the 22.55-second trial. This function used a Fourier basis set with 13 regressors. This analysis was performed separately for each participant across all regions of interest (ROIs). The ROIs were 360 cortical regions (180 in each hemisphere) (Glasser et al., 2016), 10 subcortical regions from subcortical *aseg* segmentation in FreeSurfer (Fischl et al., 2002), and hippocampal subfields extracted with the FreeSurfer hippocampal subfields module (v6.0.0; Iglesias et al., 2015). This hippocampal segmentation employs a probabilistic atlas derived from ultra-high-resolution ex vivo MRI, and was performed in multimodal mode, incorporating both T1-weighted and T2-weighted images to improve boundary definition. The cortical regions from the HCP-MMP1.0 Glasser included as MTL regions were 8, per hemisphere: (Rolls et al., 2023): EC (entorhinal cortex), PreS (presubiculum), PeEc (Perirhinal Ectorhinal), H (hippocampus), and PHA1–PHA3 (parahippocampal area 1-3), and TF (Area TF).

For each ROI, we extracted the fMRI time series, z-scored them within run, concatenated runs per participant, and then submitted them to nideconv (GroupResponseFitter) for event-related deconvolution. Event onsets were obtained from the BIDS event files. Correct, incorrect, and passive viewing trials were modeled in the deconvolution, but only correct and passive viewing trials were retained for further analysis. Following deconvolution, the extracted time courses were segmented into three task phases: encoding (0–8.25 s), delay (8.25–16.5 s), and test (16.5–22.55 s). For each phase, the mean of the time series values within the phase window was calculated for both correct and passive viewing trials. These phase-specific activity values were then used in subsequent statistical analyses.

### 2.7 Statistical Analysis

#### 2.7.1 Behavioral Analysis

All analyses were conducted in R (version 4.4.1), with visualizations created using ggplot2 (Wickham, 2016).

We assessed sex differences in accuracy, reaction time (RT), and average load level using Mann-Whitney U tests for non-normally distributed data or Welch’s t-tests for normal data with unequal variances. As no significant differences were found in accuracy or RT, and only a marginal effect on load level was observed, sex was not included as a covariate or grouping factor in the main analyses.

Memory performance above chance was assessed with a one-sample Wilcoxon signed-rank test. RT for correct and incorrect trials were compared using a Mann-Whitney U test, while the relationship between age and RT was examined via linear regression. The relationship between age and average load level was analyzed using Spearman’s rank correlation.

To assess the relationship between behavioral performance (average load level) and fMRI activation differences, we conducted correlation analyses. Normality was assessed with the Shapiro-Wilk test, which indicated that the data were not normally distributed. Therefore, we used Spearman’s rank correlation. P-values were obtained from the standard implementation in cor.test() in R, which accounts for tied ranks by providing exact or approximate significance levels.

#### 2.7.2 fMRI Data Analysis

Linear mixed-effects models were fitted using the lmerTest package in R to examine task-related activity across all participants, with event type as a fixed effect and participant as a random effect. Analyses focused on correct working-memory trials contrasted with passive viewing to isolate activity related to successful performance. As mentioned above, the passive viewing trials were matched for lower-level visual input and response requirements. This condition allowed us to isolate trial-type modulations on top of the sensory response, improving the interpretability of the observed MTL activity as participants do not attempt to perform the critical cognitive operations. Although incorrect trials have higher stimulus similarity and might better control for vigilance/effort, such trials may also be contaminated by heterogenous processes, including lapses of attention or error monitoring. Also, the number of incorrect trials, partly by design of the staircase procedure, were limited and highly variable across participants (median = 7, IQR = 4.25, range = 1–18). Specifically, 99 of 128 (77%) had <10 errors and 61 of 128 (48%) had <7 errors, which reduces the stability of condition-specific estimates, favouring the use of the passive viewing trials as control conditions (10 trials for all participants).

For group comparisons, group was included as a fixed effect. In the model comparing younger and older adults, relative motion (RMS) was also included as a covariate to account for age-related differences in head motion. To assess statistical significance, p-values for fixed effects were computed using Satterthwaite’s approximation, which is automatically applied in lmerTest to estimate denominator degrees of freedom based on variance components, providing an approximate t-distribution (Kuznetsova et al., 2017). For visualization, we combined p-values with the directionality of the effect by multiplying each p-value with the sign of the corresponding t-value, resulting in signed p-values. All brain visualizations were created using ggseg (Mowinckel & Vidal-Piñeiro, 2020).

To evaluate the appropriateness of the mixed-effects model, we examined singular fit warnings, which indicated that the random intercept variance was close to zero and did not contribute meaningfully to the model. This issue arose in approximately one-fourth of the ROIs across all phases, suggesting that the fixed effects (event type, group, and their interaction) explained nearly all variability, leaving insufficient between-subject variance to estimate the random intercept reliably. To further investigate this, we conducted separate linear models (without random effects) for these ROIs. The results showed that for ROIs with singular fit warnings, the p-values for the interaction term were identical between the mixed model and the simpler linear model, whereas ROIs without singular warnings exhibited slight variations in p-values. Since the mixed model produced the same results as the simpler models in these cases, we retained the values from the mixed model for further visualization.

To control for multiple comparisons in the subcortical analyses, we used a false discovery rate (FDR) correction (Benjamini & Hochberg, 1995). For the group analyses of the cortical regions, we applied a cluster-based permutation correction (Voldsbekk et al., 2022). Here, we assumed that engagement patterns likely would span adjacent ROIs. This method leveraged an atlas-derived adjacency matrix, where cortical ROIs were defined as neighbors if they shared a border on the HCP MMP1.0 surface (Glasser et al., 2016). Group comparisons were conducted separately for higher-performing younger adults vs. lower-performing younger adults, and lower-performing younger adults vs. older adults, with the permutation-based approach applied to identify significant clusters corrected for multiple comparisons. The clustering procedure was guided by a predefined threshold of p < .05, and cluster significance was assessed using 5000 permutations. The maximum cluster sizes from the permuted data were used to correct for multiple comparisons, yielding a corrected significance threshold. In addition to the group analyses, we conducted complementary analyses where we entered either average load level or age as a continuous predictor, to examine whether findings were consistent when the dependent variable was modeled continuously.

To interpret our results, in two instances we used Neurosynth (Yarkoni et al., 2011). First, identify established working memory regions, the Neurosynth working memory association map was thresholded at z > 10 to isolate the most strongly associated clusters, which were then cross-referenced with the HCP-MMP1.0 Glasser atlas in volumetric space to determine anatomical labels. This approach identified significant activation in left p9-46v (posterior area 9-46v, a central portion of the dorsolateral prefrontal cortex) and left AIP (Anterior Intraparietal Area), both of which are well-established regions implicated in working memory processing (Glasser et al., 2016). Second, to interpret group differences between higher and lower-performing younger adults, we employed Neurosynth to assess how regions showing differences mapped onto various cognitive functions.

## 3. Results

### 3.1 Behavioral Performance

Across all participants, memory performance was significantly above chance (defined as 33% accuracy based on three response options; V = 8256, p < .001) with no indication of a ceiling effect. Median accuracy was 76.7% (IQR = 14.2%), ranging from 40% to 96.7%. Reaction times (RT) were significantly faster for correct trials (median = 1675 ms, IQR = 1720 ms) compared to incorrect trials (median = 2235 ms, IQR = 2356 ms; W = 1369924, p < .001). There was a relationship between RT and age (b = 16 ms/year, SE = 2.5, t = 6.40, p < .001), and between age and average load level—the average number of objects participants were required to maintain in memory during the task (ρ=−0.57, p-value < .001) (**Fig. 2B**), indicating that at higher ages, participants tended to perform at lower load levels. As partly expected, given the staircase structure of the task, average load level was highly associated with accuracy, with higher load levels predicting better accuracy even after controlling for age (b = 0.105, p < .001) (**Fig. 2C**), an association that is partly expected given the staircase structure of the task.

**Figure 2.**
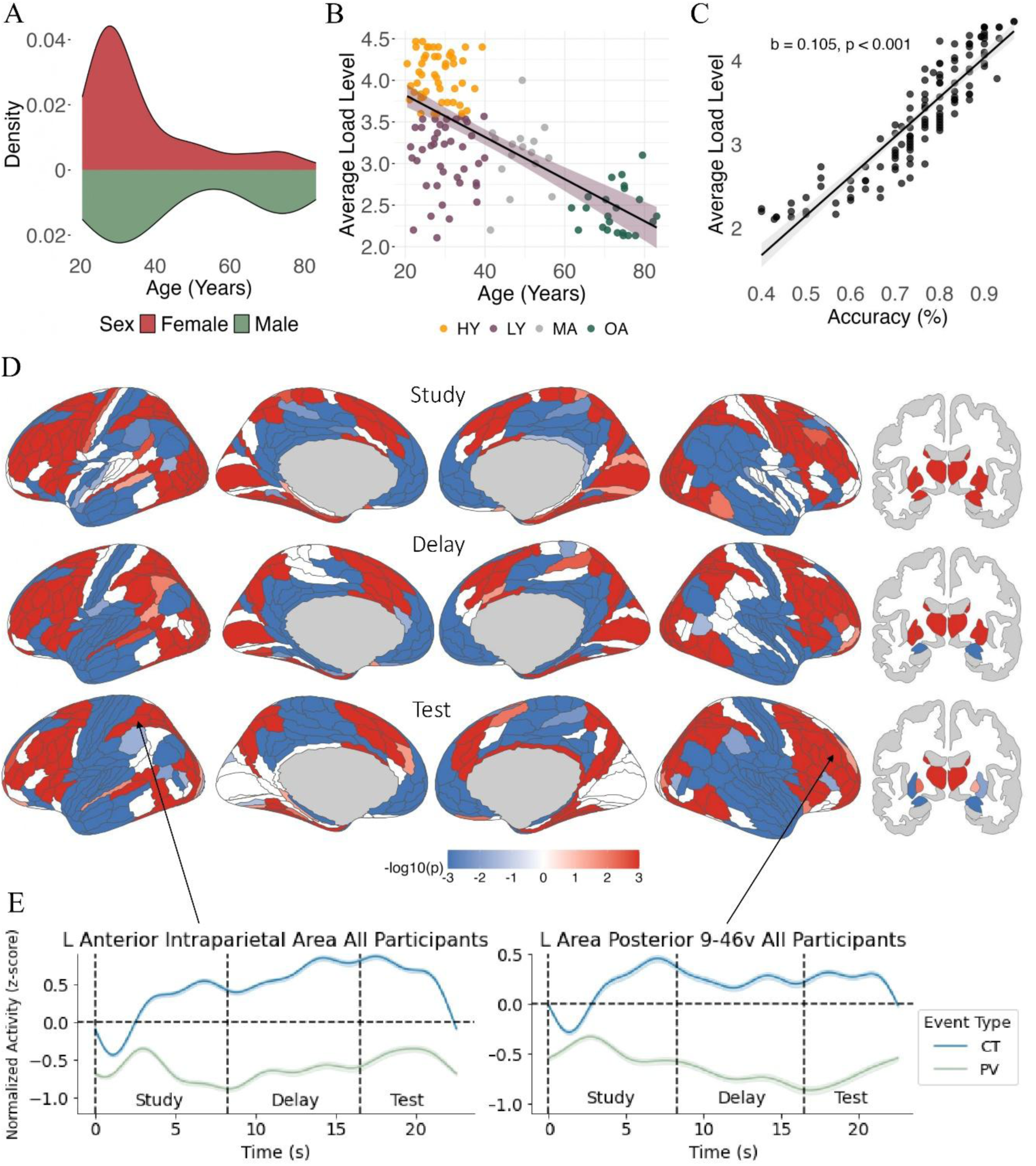
(A) Age distribution. (B) Age and average load level. Each point represents an individual participant, with the solid black line indicating the linear regression fit. The shaded area represents the 95% confidence interval for the regression line. OA = Older Adults, MA = Middle-Aged, HY = Higher-Performing Younger Adults, LY = Lower-Performing Younger Adults. (C) Accuracy and average load level. (D) Activity differences in the contrast between correct trials vs. passive viewing (FDR corrected for multiple comparisons), shown as log-transformed p-values. Red areas indicate regions where correct trials elicited greater activity than passive viewing, while blue areas indicate regions where correct trials showed lower activity than passive viewing. Regions with P_FDR_ > .05 in white. (E) Trial-averaged time series for both correct trials and passive viewing conditions in well-established working memory regions, left AIP (left) and left p9-46v (right). Shaded areas show standard error across participants. Arrows indicate the respective ROIs. *Note:* The arrow for area p9-46v is placed on the right hemisphere for visual clarity, as positioning it on the left hemisphere would obstruct key plot elements. **Abbreviations:** P_FDR_ = false discovery rate-corrected p-value; AIP = anterior intraparietal area; p9-46v = posterior area 9-46v

### 3.2 Task-Related Activation in Established Working Memory Regions

To contextualize our findings within established working memory activity patterns, we first looked at brain activity patterns during correct performance of the task (**Fig. 2D**). Across all participants, we compared task activation during correct trials with activation during passive viewing. Most cortical regions (85.9%) showed differences (PFDR range: 3.72×10^-70^ to .047), with frontoparietal areas showing greater activation during correct trials. Conversely, regions such as the medial prefrontal cortex and posterior cingulate cortex exhibited greater activation during passive viewing. Both p9-46v and AIP exhibited sustained activation throughout the trial compared to passive viewing (**Fig. 2E**), consistent with their well-established role in working memory (Rottschy et al., 2012). Having obtained support for the involvement of these canonical working memory regions, we next examined the role of medial temporal lobe (MTL) regions in this allocentric spatial working memory task.

### 3.3 Activity During Spatial Working Memory in MTL

#### 3.3.1 Hippocampal and Subcortical Activity During Correct Trials Across Participants

Following an approach similar to Hannula and Ranganath (2008), we examined hippocampal activity across the study, delay, and test phases. Based on all participants, **Fig. 3A** shows examples of activity across the event phases from left anterior and posterior hippocampus. Using mixed models separately in each of the three event phases, we tested differences in activity between event types, that is, between correct trials and passive viewing. As shown in **Fig. 3B**, nearly all hippocampal subregions exhibited a significant effect of event type across task phases, reflecting either active task engagement or task-induced deactivation, depending on the phase. T- and p-values ranged from t(127) = -9.86, PFDR = 4.26×10^-16^ to t(254) = -1.66, PFDR = .016. The only non-significant results were observed in the left and right posterior hippocampus during the delay phase (t(127) = -1.66, PFDR = .10 and t(127) = -0.76, PFDR = .45 respectively). Besides the posterior hippocampus also differed by displaying higher activation again during the test phase, all hippocampal subregions demonstrated greater activation during the study phase compared to passive viewing, whereas the delay and test phases were characterized by greater deactivation.

**Figure 3.**
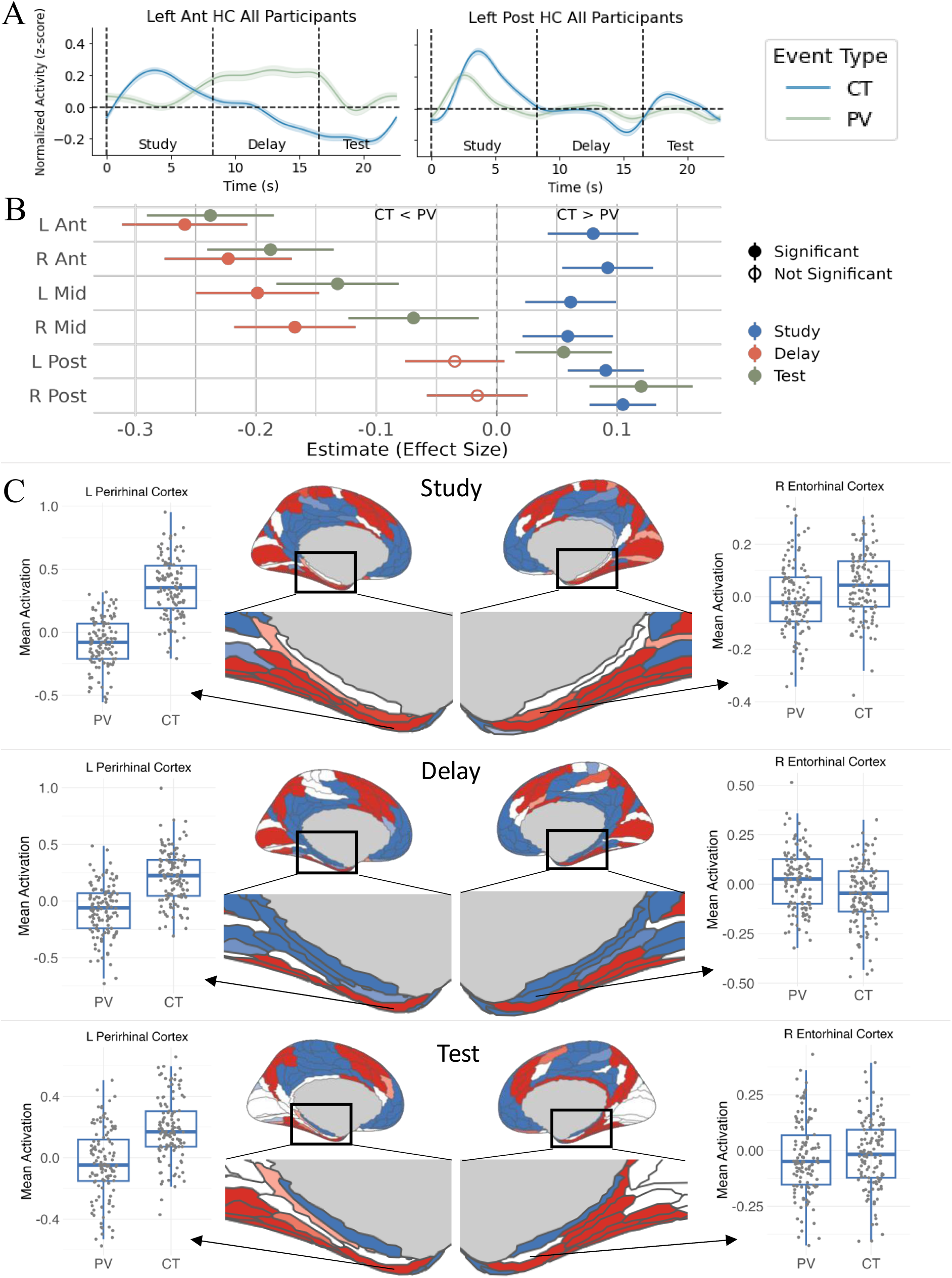
(A) Trial-averaged time series of left anterior and posterior hippocampus across all participants. Shaded areas show standard error. (B) The hippocampal subregions in different phases. The x-axis shows the estimated difference in normalized brain activity (z-scores) between the mean activity of correct trials and passive viewing for each phase. Positive and negative values reflect greater activity during correct trials and passive viewing, respectively. Points represent the estimated effect, and horizontal lines indicate the 95% confidence intervals. (C) Cortical MTL activation patterns. Red and blue areas indicate greater activity in correct trials and passive viewing, respectively. Regions with P_FDR_ > .05 in white. Boxplots for the left perirhinal ectorhinal cortex and right entorhinal cortex are included for all three phases. **Abbreviations:** P_FDR_ = false discovery rate-corrected p-value; MTL = medial temporal lobe.

Beyond the hippocampus, all other subcortical regions also exhibited significant differences between correct trials and passive viewing across all three phases, with t and p values ranging from t(127) = 16.50, PFDR = 7.37×10^-32^ in the right putamen during delay phase to t(127) = 2.00, PFDR = .047 in the right pallidum during test phase (**Fig. 2D**).

#### 3.3.2 Cortical MTL Activity Across Participants

To examine cortical MTL regions, we revisited the contrast between correct trials and passive viewing presented above. Significant effects were observed in 41of 48 MTL ROI-by-phase tests (PFDR range: 3.69×10⁻³⁹ to .045), mirroring the phase-specific patterns seen in the hippocampus. Most cortical MTL regions exhibited greater activity during correct trials compared with passive viewing in the study phase (**Fig. 3C**). During the delay phase, distinct patterns of activity emerged across regions. Similar to the hippocampus, the bilateral entorhinal cortex, presubiculum, and parahippocampal area 1 showed lower activity during correct trials than passive viewing. In contrast, regions such as the bilateral perirhinal ectorhinal cortex, parahippocampal area 3, and right parahippocampal area 2 displayed the opposite pattern, with higher activity during correct trials (see boxplots of the perirhinal cortex in **Fig. 3C**). In the test phase, several MTL regions again exhibited increased activation during correct trials, alongside the bilateral posterior hippocampus, except for bilateral entorhinal cortex and left parahippocampal area 1 which no longer exhibited any difference. Overall, MTL regions, including both the hippocampus and cortical areas, showed greater activation during correct trials than passive viewing in the study phase, widespread deactivation during the delay phase with some regions showing sustained activity, and a return to increased activation in the test phase, although anterior and mid portions of the hippocampus remained deactivated, and several areas no longer exhibited differences.

### 3.4 Activity Difference between Performance- and Age-based Groups

#### 3.4.1 Activity Differences Between Higher and Lower-Performing Younger Adults

As exemplified in **Fig. 4A** for left anterior hippocampus, mixed models of group differences in the differences between correct trials and passive viewing showed no differences between higher- and lower-performing younger adults in the hippocampus or other subcortical structures (lowest p-value = .097, the left posterior hippocampus, delay phase).

**Figure 4.**
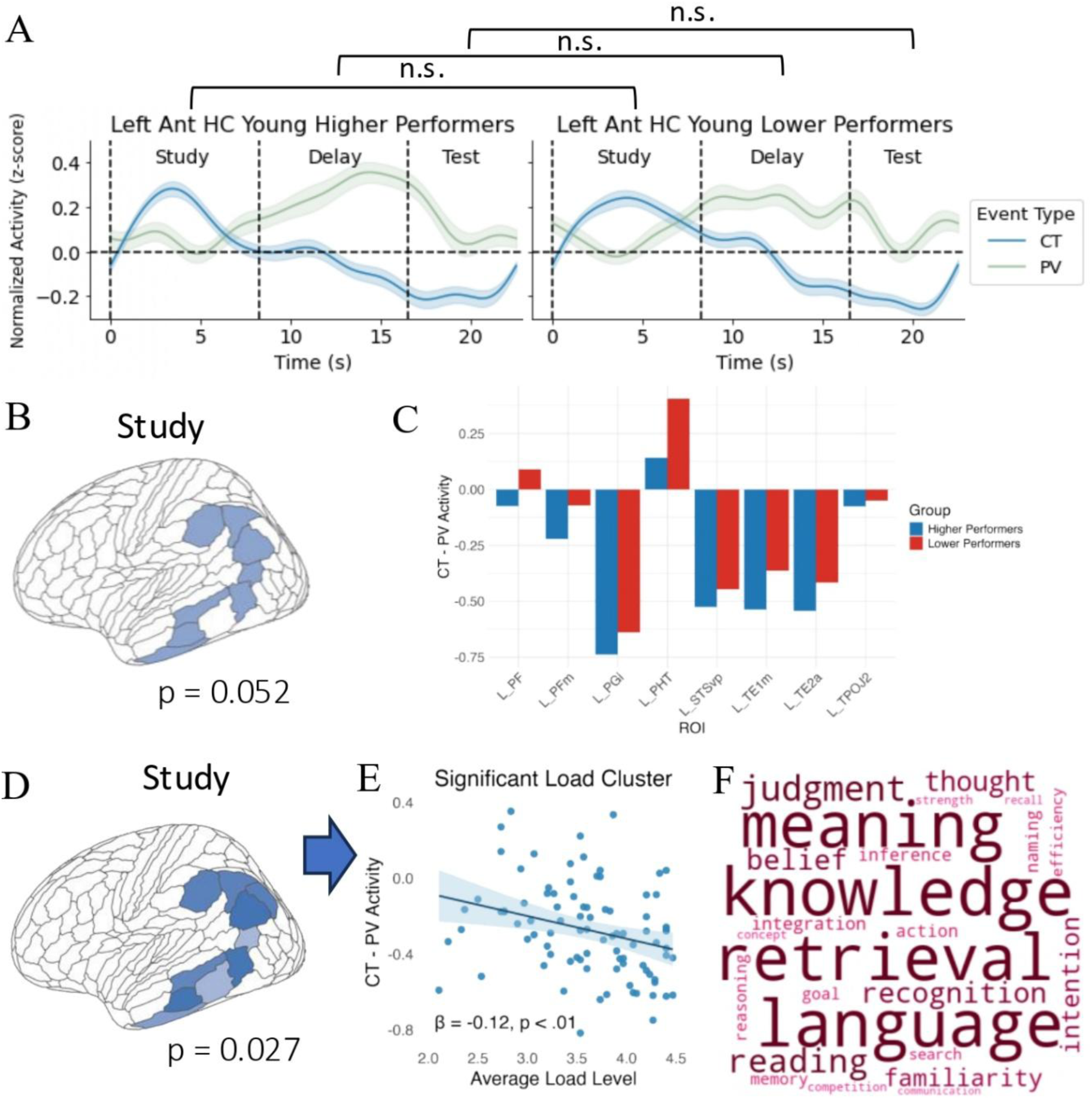
(A) Trial-averaged time series of activation in the left posterior hippocampus for higher- and lower-performing younger adults, respectively. Shaded areas show standard error across participants. (B) Group analyses cluster in the left hemisphere during the study phase demonstrating a weak interaction (P_perm_ = .052). (C) Activation in the eight cluster regions. (D) Cluster reflecting the effect of individuals’ average load level as a fixed factor (P_perm_ = .027). (E) The association between average load level and correct trials–passive viewing difference extracted from the cluster in D. (F) Word cloud of terms most strongly associated with the observed cluster. **Abbreviations**: n.s. = not significant; P_FDR_ = false discovery rate-corrected p-value; CT = correct trials; PV = passive viewing.

Cortically, across phases, 15 clusters were initially identified at the cluster-forming threshold of p < .05 uncorrected. However, none survived correction with the permutation test (all Pperm ≥ .07), with the lowest corrected p-value observed in the test phase. However, one cluster in the left hemisphere including eight regions (**Fig. 4B**) showed a strong trend towards an interaction during the study phase (Pperm = .052). Here, a numerically stronger deactivation for higher performers is observed in correct trials vs passive viewing in most of the regions in the cluster (**Fig. 4C**). Modeling performance as a continuous predictor (see Methods), revealed a stronger association for a similar cluster including all the same regions (Pperm = .027) (**Fig. 4D**). Here, higher average load level was likewise associated with stronger deactivation in correct trials compared to passive viewing during the study phase (**Fig. 4E**). The areas showing differences were primarily associated with language, meaning, and knowledge (**Fig. 4F**).

As our focus is on MTL involvement, we conducted an additional sensitivity analysis to ensure that no significant correlations were overlooked. Specifically, we tested the relationship between activation in MTL regions and task performance. However, our analysis did not reveal a correlation in any MTL regions, with p-values ranging from PFDR = .71 to PFDR = .98, including the left entorhinal cortex during the delay phase found in Hannula and Ranganath (2008), reinforcing our initial findings of no significant MTL involvement in performance differences among young adults.

#### 3.4.2 Activity Differences Between Older Adults and Lower-Performing Younger Adults

To investigate age-related differences, we compared activation patterns between older adults and the lower-performing younger adults, which showed the most comparable performance level, although still significantly different (2.88 vs 2.44 for lower-performing younger adults and older adults, respectively, p < .001). As shown in **Fig. 5A**, older adults exhibited a smaller difference between correct trials and passive viewing during the test phase in the anterior hippocampus, reflecting a significant event × group interaction in both the left (t(131) = 3.70, PFDR = .006), and right anterior hippocampus (t(131) = 3.39, PFDR = .008). The remaining hippocampal tests did not show any group differences (PFDR > .21).

**Figure 5.**
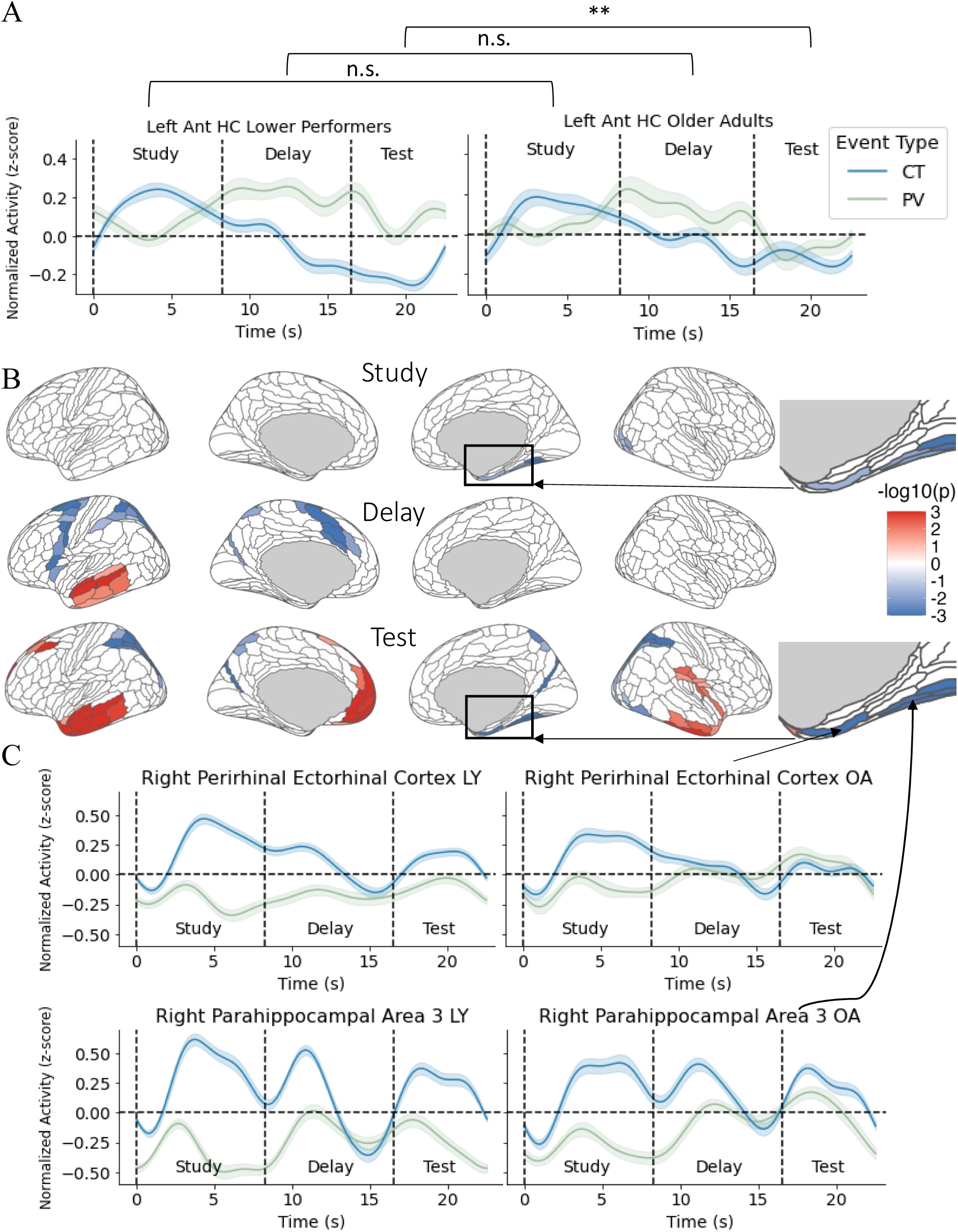
(A) Trial-averaged time series of left anterior hippocampus in lower-performing younger adults and older adults resulting in significant group difference in the test phase. (B) Cortical regions showing interaction effects, shown as log-transformed, cluster-corrected p-values. Across all clusters, lower-performing younger adults exhibited a greater difference between correct trials and passive viewing than older adults. Blue areas show regions where correct trials–passive viewing values were lower in older adults while red areas show regions where correct trials-passive viewing values were higher in older adults compared to younger adults. However, this reflects reduced overall activity rather than a larger difference between correct trials and passive viewing. (C) Trial-averaged time series of right perirhinal ectorhinal cortex and right parahippocampal area 3 in lower-performing younger adults and older adults resulting in significant group difference in the study and test phases values. Shaded areas show standard error. **Abbreviations**: n.s. = not significant; CT = correct trials; PV = passive viewing.

Further group comparisons revealed differences between lower-performing younger adults and older adults across both subcortical and cortical regions. In the subcortical analysis, the mixed-effects model identified a significantly greater activation difference between correct trials and passive viewing in older adults compared to lower-performing younger adults in the left putamen (t(70) = 3.72, PFDR = .012) during the test phase. Cortical analysis **(Fig. 5B)** revealed significant interactions between event type and group in 9 significant clusters across all three phases. MTL regions – the right perirhinal ectorhinal cortex and parahippocampal area 3 – were observed in significant clusters in the study (PPERM = .026) and test (PPERM = .004) phases. As shown in **Fig. 5C**, in the significant MTL regions, lower-performing younger adults exhibited a larger correct trials–passive viewing difference than older adults, with passive viewing values remaining around zero and correct trials showing activation. Modeling age as a continuous predictor did not alter the MTL findings; however, additional clusters emerged in the left prefrontal cortex during the delay phase and in the left temporal lobe during the test phase.

Together, these results indicate that age-related differences in task-related activity extend beyond the hippocampus into surrounding MTL regions. The findings suggest that older adults generally exhibit reduced correct trials–passive viewing differences, with some variations across specific MTL regions.

## 4. Discussion

We investigated the role of the hippocampus and cortical areas in allocentric working memory. Our findings support hippocampal and cortical MTL involvement in encoding, maintenance, and retrieval, with the hippocampus showing a distinct anterior-posterior gradient across phases. However, contrary to the activity in established working memory regions like the lateral prefrontal cortex, this hippocampal and cortical MTL engagement did not manifest as sustained BOLD activity, although perirhinal and parahippocampal cortices were activated in all three phases. Notably, activation differences between higher- and lower-performing younger adults were only observed in a left temporoparietal cluster at encoding. Contrasting lower-performing younger adults with older adults, however, showed test phase activation differences in the anterior hippocampus. Age-related differences were also observed in right perirhinal and parahippocampal cortices, where activation showed lower differentiation between correct and control trials in older adults. These cortical differences were present during the study and test phase but not the delay phase. Taken together, these findings demonstrate hippocampal and cortical MTL involvement across the temporal unfolding of an allocentric working memory task requiring complex relational processing, including the delay phase. That performance-related differences among younger adults emerged in a left temporoparietal cluster rather than in MTL regions, however, suggests that this engagement, while robust, may play a more auxiliary role.

The hippocampus exhibited modulations of activity during all phases of a spatial working memory task, providing support for its involvement throughout the temporal unfolding of allocentric short-term processing. In line with hippocampal engagement in encoding spatial representations (Parslow et al., 2004; Suthana et al., 2009), study activation in anterior, mid and posterior subfields were all higher during correct trials compared with passive viewing. This widespread involvement mirrors the results of Hannula and Ranganath (2008) and aligns with a meta-analysis by Sullivan et al. (2024) showing reliable recruitment of both anterior and posterior hippocampus during spatial encoding.

During the delay phase, anterior and mid hippocampal portions showed deactivation for correct trials compared with passive viewing. This deactivation contrasts with the sustained activation pattern, of magnitudes greater than zero, observed for intraparietal and lateral prefrontal cortices (**Fig. 2E**), and, in the MTL, parahippocampal and perirhinal (**Fig. 5C, 3C**) but not entorhinal cortices (**Fig. 3C**), all putatively involved in working memory maintenance (Christophel et al., 2017; Schon et al., 2016). Reviewing 26 fMRI studies, Slotnick (2022) contends that working memory does not activate the hippocampus, as it fails to show sustained activation during the late delay period – the only phase assumed to isolate working memory maintenance from long-term memory processes. Consistent with this perspective, we did not observe positive BOLD signals in the hippocampus during the delay phase. Neither did we observe performance- or age-related differences in hippocampal delay signal. Yet, although BOLD decreases are often interpreted as reduced involvement, this view has been challenged. For example, fMRI shows hippocampal deactivation during relational attention alongside stable multivariate representations (Córdova et al., 2019). More broadly, delay information can be carried by transient bursts, dynamic population codes, or sparse assemblies that need not yield sustained BOLD (Sreenivasan & D’Esposito, 2019). Together with the complexity of neurovascular coupling in the hippocampus (Ekstrom, 2010), the observed hippocampal deactivation may reflect a shift in processing mode rather than disengagement.

If the hippocampus played no role during the delay, we would expect the activity to resemble passive viewing. Instead, only the posterior hippocampus showed this pattern, while the anterior and mid regions engagement differed between correct trials and passive viewing. Because passive viewing trials consisted of one familiar object, hippocampal passive viewing activity cannot readily be explained as extended encoding of novel information in the early delay phase (Slotnick, 2022), which has been linked with long-term memory encoding (Nauer et al., 2015; Ranganath et al., 2005). Although we did not model early and late delay separately, visual inspection of the curves did not point to early delay processes driving the overall delay pattern on correct or passive viewing trials (Fig. 3A).

Summarizing the delay phase hippocampal results, we cannot rule out that the deactivation reflects lack of maintenance engagement. Still, the observed pattern — hippocampal deactivation differing from passive viewing, alongside positive parahippocampal and perirhinal cortical activation — suggests that both hippocampal and cortical MTL regions contribute during the delay, though potentially through different mechanisms. The results are thus compatible with anterior and mid subregions supporting delay processes via mechanisms not captured by sustained BOLD signal alone. Such a hippocampal support role accords with multivariate fMRI evidence (Libby et al., 2014) and intracranial electrophysiological recordings of hippocampal delay maintenance (Kamiński et al., 2017; Peters & Reithler, 2022). The overall pattern, that is, hippocampal engagement most evident during encoding and retrieval, with cortical MTL regions showing positive activation during the delay, is broadly consistent with the interaction between representational complexity and retention interval proposed by Yonelinas (2013), although the continued differentiation of hippocampal delay activity from passive viewing suggests that the boundary between hippocampal and cortical contributions is less categorical than a strict division by task phase would imply.

During the test phase, posterior hippocampal activation re-emerged alongside activation in several cortical MTL regions, mirroring the results of Hannula and Ranganath (2008). This posterior hippocampal reactivation, in contrast to the continued deactivation of anterior and mid subregions, points to distinct functional contributions along the hippocampal long axis across task phases.

Cortical MTL involvement was observed across most regions, including entorhinal, perirhinal ectorhinal, presubiculum, and parahippocampal cortices, with greater activation for correct trials than passive viewing in both study and test phases. In contrast, the delay phase showed widespread deactivation, particularly in the entorhinal cortex, parahippocampal area 1, and presubiculum. However, bilateral perirhinal ectorhinal cortex, parahippocampal area 3, and right parahippocampal area 2 showed greater activation for correct trials than passive viewing in all three phases. This result partly aligns with Schon et al. (2016), who observed sustained delay activity in entorhinal, perirhinal, and parahippocampal cortices during a scene-matching task, interpreted as support for a working memory buffer in these regions. In our data, bilateral perirhinal and parahippocampal cortices similarly showed greater delay-phase activation for correct trials, while the entorhinal cortex showed deactivation. Whereas the perirhinal cortex is implicated in familiarity and object representation, the parahippocampal cortex has been linked to spatial and contextual processing, including scene representation and environmental layout (Fiorilli et al., 2021; Ranganath & Ritchey, 2012).

Contrary to our second hypothesis, we did not find different MTL activation in higher-performing younger adults. Hence, we did not replicate the finding of Hannula and Ranganath (2008) of an association between performance and early-delay activity in the entorhinal cortex. Instead, performance was associated with stronger modulation of the correct–passive viewing control contrast during the study phase in a left temporoparietal cluster. Given that these regions have been linked to language-related processes (Price, 2012), the observed effect may reflect the use of verbal strategies to support relational encoding. Speculatively, some participants may have verbalized object identities and spatial layouts during encoding, relying on linguistic scaffolding as an alternative or supplement to MTL-dependent mechanisms. However, as discussed above, there are challenges in interpreting activation and deactivation patterns with respect to neural engagement. Hence, whether such verbalization was performance beneficial (higher performers showed a larger difference between correct and control trials), or detrimental (higher performers showed more negative values), remain unclear with the current data. Future studies could manipulate or assess use of verbalization strategies.

In line with our third prediction, older adults exhibited lower hippocampal and cortical MTL activation, particularly during the test phase. Specifically, activation differences in bilateral anterior hippocampus were smaller compared with younger adults. Studies have similarly reported lower anterior hippocampus involvement during context-rich retrieval in older adults (Kukolja et al., 2009; Mitchell et al., 2000). Additionally, we observed smaller condition-related differences in the parahippocampal and perirhinal cortices. Reanalyzing data from Hannula & Ranganath (2008), Libby et al. (2014) found that that successful maintenance of object-location relations associated with stronger pattern similarity of the encoding and delay activity in the anterior hippocampus. Extending this notion of a role for anterior hippocampus in identity-location integration, lower anterior deactivation in our older group may reflect impaired retrieval of integrated representations. Similarly, smaller differences in perirhinal and parahippocampal activation may reflect reduced object and context coding, respectively (Libby et al., 2014).

Several limitations should be considered. First, our event-related design modeled study, delay, and test within a fixed trial. Without within-trial jitter or partial-trial omissions, the phase regressors are partly collinear. Deconvolution reduces but cannot eliminate phase overlap, so later-phase estimates may retain variance from previous phases. Study-phase estimates can therefore be interpreted with greater confidence than later phases. Second, we used a staircase to better match task difficulty across individuals and age groups, but the step size (±1 object) and bounds (minimum two; maximum five) limited granularity. The procedure targeted 70.7% accuracy, yet some participants remained above or below this level. Third, the staircase design yields more equity in effort or performance, but achieved load still differed between participants and groups, illustrating the inherent difficulty of obtaining equitable comparisons in this type of task. Fourth, due to a restricted number of trials, we did not assess trials sorted by trial type (match, mismatch-position, mismatch-swap), which may influence activity particularly in the test phase.

In conclusion, these findings suggest the involvement of hippocampal and cortical MTL subregions throughout all phases of allocentric working memory, with distinct roles emerging across the temporal unfolding of the task. One approach for future studies could be an experimental design with within-trial jitter or partial-trial omission to further reduce phase estimate overlap. Another intriguing approach would be to use focused transcranial ultrasound stimulation, a non-invasive technique able to modulate deep brain structures (Murphy et al., 2025), to test causal effects of hippocampal disruption on allocentric working memory at various stages.

## Data and Code Availability

The data and analysis code supporting the findings of this study can be made available from the corresponding author upon publication. Raw MRI data are not publicly available due to ethical/legal restrictions related to participant privacy.

## Author Contributions

All authors designed the research. Sneve performed preprocessing. Orvik, Grydeland, and Sneve analysed the data. Orvik and Grydeland drafted the manuscript, and all authors reviewed, edited, and approved the final manuscript.

## Declaration of Competing Interests

The authors declare no competing financial or non-financial interests.

## Acknowledgments

This work was supported by the Research Council of Norway [grant #325415 to H.G].

## Notes

### Competing Interest Statement

The authors have declared no competing interest.

### Summary of Updates

This revised version updates the manuscript framing to better situate the study within the broader literature on hippocampal and medial temporal lobe contributions to working memory. The introduction and discussion have been revised to clarify the relationship between the present task, relational memory, and previous work beyond the original Hannula and Ranganath framework.

## References

Andersson, J. L. R., Skare, S., & Ashburner, J. (2003). How to correct susceptibility distortions in spin-echo echo-planar images: Application to diffusion tensor imaging. NeuroImage, 20(2), 870–888. 10.1016/S1053-8119(03)00336-7

Benjamini, Y., & Hochberg, Y. (1995). Controlling the False Discovery Rate: A Practical and Powerful Approach to Multiple Testing. Journal of the Royal Statistical Society Series B: Statistical Methodology, 57(1), 289–300. 10.1111/j.2517-6161.1995.tb02031.x

Burgess, N. (2008). Spatial Cognition and the Brain. Annals of the New York Academy of Sciences, 1124(1), 77–97. 10.1196/annals.1440.002

Christophel, T. B., Klink, P. C., Spitzer, B., Roelfsema, P. R., & Haynes, J.-D. (2017). The Distributed Nature of Working Memory. Trends in Cognitive Sciences, 21(2), 111–124. 10.1016/j.tics.2016.12.007

Córdova, N. I., Turk-Browne, N. B., & Aly, M. (2019). Focusing on what matters: Modulation of the human hippocampus by relational attention. Hippocampus, 29(11), 1025– 1037. 10.1002/hipo.23082

Ekstrom, A. (2010). How and when the fMRI BOLD signal relates to underlying neural activity: The danger in dissociation. Brain Research Reviews, 62(2), 233–244. 10.1016/j.brainresrev.2009.12.004

Fiorilli, J., Bos, J. J., Grande, X., Lim, J., Düzel, E., & Pennartz, C. M. A. (2021). Reconciling the object and spatial processing views of the perirhinal cortex through task-relevant unitization. Hippocampus, 31(7), 737–755. 10.1002/hipo.23304

Fischl, B. (2012). FreeSurfer. NeuroImage, 62(2), 774–781. 10.1016/j.neuroimage.2012.01.021

Fischl, B., Salat, D. H., Busa, E., Albert, M., Dieterich, M., Haselgrove, C., Van Der Kouwe, A., Killiany, R., Kennedy, D., Klaveness, S., Montillo, A., Makris, N., Rosen, B., & Dale, A. M. (2002). Whole Brain Segmentation. Neuron, 33(3), 341–355. 10.1016/S0896-6273(02)00569-X

Folstein, M. F., Folstein, S. E., & McHugh, P. R. (1975). “Mini-mental state”. Journal of Psychiatric Research, 12(3), 189–198. 10.1016/0022-3956(75)90026-6

Frederick, B. deB., Nickerson, L. D., & Tong, Y. (2012). Physiological denoising of BOLD fMRI data using Regressor Interpolation at Progressive Time Delays (RIPTiDe) processing of concurrent fMRI and near-infrared spectroscopy (NIRS). NeuroImage, 60(3), 1913–1923. 10.1016/j.neuroimage.2012.01.140

Glasser, M. F., Coalson, T. S., Robinson, E. C., Hacker, C. D., Harwell, J., Yacoub, E., Ugurbil, K., Andersson, J., Beckmann, C. F., Jenkinson, M., Smith, S. M., & Van Essen, D. C. (2016). A multi-modal parcellation of human cerebral cortex. Nature, 536(7615), 171–178. 10.1038/nature18933

Glasser, M. F., Sotiropoulos, S. N., Wilson, J. A., Coalson, T. S., Fischl, B., Andersson, J. L., Xu, J., Jbabdi, S., Webster, M., Polimeni, J. R., Van Essen, D. C., & Jenkinson, M. (2013). The minimal preprocessing pipelines for the Human Connectome Project. NeuroImage, 80, 105–124. 10.1016/j.neuroimage.2013.04.127

Grady, C. L. (2020). Meta-analytic and functional connectivity evidence from functional magnetic resonance imaging for an anterior to posterior gradient of function along the hippocampal axis. Hippocampus, 30(5), 456–471. 10.1002/hipo.23164

Griffanti, L., Salimi-Khorshidi, G., Beckmann, C. F., Auerbach, E. J., Douaud, G., Sexton, C. E., Zsoldos, E., Ebmeier, K. P., Filippini, N., Mackay, C. E., Moeller, S., Xu, J., Yacoub, E., Baselli, G., Ugurbil, K., Miller, K. L., & Smith, S. M. (2014). ICA-based artefact removal and accelerated fMRI acquisition for improved resting state network imaging. NeuroImage, 95, 232–247. 10.1016/j.neuroimage.2014.03.034

Hannula, D. E., & Ranganath, C. (2008). Medial Temporal Lobe Activity Predicts Successful Relational Memory Binding. The Journal of Neuroscience, 28(1), 116–124. 10.1523/JNEUROSCI.3086-07.2008

Hartley, T., Trinkler, I., & Burgess, N. (2004). Geometric determinants of human spatial memory. Cognition, 94(1), 39–75. 10.1016/j.cognition.2003.12.001

Hollander, G. D., Knapen, T., Eort, & Snoek, L. (2019). VU-Cog-Sci/nideconv: Release candidate 4 (Version 0.1-rc4) [Computer software]. Zenodo. 10.5281/ZENODO.1463838

Idland, A.-V., Sala-Llonch, R., Borza, T., Watne, L. O., Wyller, T. B., Brækhus, A., Zetterberg, H., Blennow, K., Walhovd, K. B., & Fjell, A. M. (2017). CSF neurofilament light levels predict hippocampal atrophy in cognitively healthy older adults. Neurobiology of Aging, 49, 138–144. 10.1016/j.neurobiolaging.2016.09.012

Iglesias, J. E., Augustinack, J. C., Nguyen, K., Player, C. M., Player, A., Wright, M., Roy, N., Frosch, M. P., McKee, A. C., Wald, L. L., Fischl, B., & Van Leemput, K. (2015). A computational atlas of the hippocampal formation using ex vivo, ultra-high resolution MRI: Application to adaptive segmentation of in vivo MRI. NeuroImage, 115, 117–137. 10.1016/j.neuroimage.2015.04.042

Jeneson, A., & Squire, L. R. (2012). Working memory, long-term memory, and medial temporal lobe function. Learning & Memory, 19(1), 15–25. 10.1101/lm.024018.111

Kamiński, J., Sullivan, S., Chung, J. M., Ross, I. B., Mamelak, A. N., & Rutishauser, U. (2017). Persistently active neurons in human medial frontal and medial temporal lobe support working memory. Nature Neuroscience, 20(4), 590–601. 10.1038/nn.4509

Korponay, C., Janes, A. C., & Frederick, B. B. (2024). Brain-wide functional connectivity artifactually inflates throughout functional magnetic resonance imaging scans. Nature Human Behaviour, 8(8), 1568–1580. 10.1038/s41562-024-01908-6

Kukolja, J., Thiel, C. M., Wilms, M., Mirzazade, S., & Fink, G. R. (2009). Ageing-related changes of neural activity associated with spatial contextual memory. Neurobiology of Aging, 30(4), 630–645. 10.1016/j.neurobiolaging.2007.08.015

Kuznetsova, A., Brockhoff, P. B., & Christensen, R. H. B. (2017). lmerTest Package: Tests in Linear Mixed Effects Models. Journal of Statistical Software, 82(13), 1–26. 10.18637/jss.v082.i13

Libby, L. A., Hannula, D. E., & Ranganath, C. (2014). Medial Temporal Lobe Coding of Item and Spatial Information during Relational Binding in Working Memory. The Journal of Neuroscience, 34(43), 14233–14242. 10.1523/JNEUROSCI.0655-14.2014

Mitchell, K. J., Johnson, M. K., Raye, C. L., & D’Esposito, M. (2000). fMRI evidence of age-related hippocampal dysfunction in feature binding in working memory. Cognitive Brain Research, 10(1–2), 197–206. 10.1016/S0926-6410(00)00029-X

Mowinckel, A. M., & Vidal-Piñeiro, D. (2020). Visualization of Brain Statistics With R Packages *ggseg* and *ggseg3d*. Advances in Methods and Practices in Psychological Science, 3(4), 466–483. 10.1177/2515245920928009

Murphy, K. R., Nandi, T., Kop, B., Osada, T., Lueckel, M., N’Djin, W. A., Caulfield, K. A., Fomenko, A., Siebner, H. R., Ugawa, Y., Verhagen, L., Bestmann, S., Martin, E., Butts Pauly, K., Fouragnan, E., & Bergmann, T. O. (2025). A practical guide to transcranial ultrasonic stimulation from the IFCN-endorsed ITRUSST consortium. Clinical Neurophysiology, 171, 192–226. 10.1016/j.clinph.2025.01.004

Nauer, R. K., Whiteman, A. S., Dunne, M. F., Stern, C. E., & Schon, K. (2015). Hippocampal subfield and medial temporal cortical persistent activity during working memory reflects ongoing encoding. Frontiers in Systems Neuroscience, 9. 10.3389/fnsys.2015.00030

Nyberg, L., & Pudas, S. (2019). Successful Memory Aging. Annual Review of Psychology, 70(1), 219–243. 10.1146/annurev-psych-010418-103052

Olson, I. R., Page, K., Moore, K. S., Chatterjee, A., & Verfaellie, M. (2006). Working Memory for Conjunctions Relies on the Medial Temporal Lobe. The Journal of Neuroscience, 26(17), 4596–4601. 10.1523/JNEUROSCI.1923-05.2006

Parslow, D. M., Rose, D., Brooks, B., Fleminger, S., Gray, J. A., Giampietro, V., Brammer, M. J., Williams, S., Gasston, D., Andrew, C., Vythelingum, G. N., Loannou, G., Simmons, A., & Morris, R. G. (2004). Allocentric Spatial Memory Activation of the Hippocampal Formation Measured With fMRI. Neuropsychology, 18(3), 450–461. 10.1037/0894-4105.18.3.450

Peters, J. C., & Reithler, J. (2022). Hippocampal activity in working memory tasks: Sparse, yet relevant. Cognitive Neuroscience, 13(3–4), 212–214. 10.1080/17588928.2022.2131746

Price, C. J. (2012). A review and synthesis of the first 20years of PET and fMRI studies of heard speech, spoken language and reading. NeuroImage, 62(2), 816–847. 10.1016/j.neuroimage.2012.04.062

Ranganath, C., Cohen, M. X., & Brozinsky, C. J. (2005). Working Memory Maintenance Contributes to Long-term Memory Formation: Neural and Behavioral Evidence. Journal of Cognitive Neuroscience, 17(7), 994–1010. 10.1162/0898929054475118

Ranganath, C., & D’Esposito, M. (2001). Medial Temporal Lobe Activity Associated with Active Maintenance of Novel Information. Neuron, 31(5), 865–873. 10.1016/S0896-6273(01)00411-1

Ranganath, C., & Ritchey, M. (2012). Two cortical systems for memory-guided behaviour. Nature Reviews Neuroscience, 13(10), 713–726. 10.1038/nrn3338

Rolls, E. T., Wirth, S., Deco, G., Huang, C., & Feng, J. (2023). The human posterior cingulate, retrosplenial, and medial parietal cortex effective connectome, and implications for memory and navigation. Human Brain Mapping, 44(2), 629–655. 10.1002/hbm.26089

Rottschy, C., Langner, R., Dogan, I., Reetz, K., Laird, A. R., Schulz, J. B., Fox, P. T., & Eickhoff, S. B. (2012). Modelling neural correlates of working memory: A coordinate-based meta-analysis. NeuroImage, 60(1), 830–846. 10.1016/j.neuroimage.2011.11.050

Salimi-Khorshidi, G., Douaud, G., Beckmann, C. F., Glasser, M. F., Griffanti, L., & Smith, S. M. (2014). Automatic denoising of functional MRI data: Combining independent component analysis and hierarchical fusion of classifiers. NeuroImage, 90, 449–468. 10.1016/j.neuroimage.2013.11.046

Schon, K., Newmark, R. E., Ross, R. S., & Stern, C. E. (2016). A Working Memory Buffer in Parahippocampal Regions: Evidence from a Load Effect during the Delay Period. Cerebral Cortex, 26(5), 1965–1974. 10.1093/cercor/bhv013

Slotnick, S. D. (2022). Does working memory activate the hippocampus during the late delay period? Cognitive Neuroscience, 13(3–4), 182–207. 10.1080/17588928.2022.2075842

Smith, S. M., Jenkinson, M., Woolrich, M. W., Beckmann, C. F., Behrens, T. E. J., Johansen-Berg, H., Bannister, P. R., De Luca, M., Drobnjak, I., Flitney, D. E., Niazy, R. K., Saunders, J., Vickers, J., Zhang, Y., De Stefano, N., Brady, J. M., & Matthews, P. M. (2004). Advances in functional and structural MR image analysis and implementation as FSL. NeuroImage, 23, S208–S219.\ 10.1016/j.neuroimage.2004.07.051

Sreenivasan, K. K., & D’Esposito, M. (2019). The what, where and how of delay activity. Nature Reviews Neuroscience, 20(8), 466–481. 10.1038/s41583-019-0176-7

Sullivan, M. A., Fritch, H. A., & Slotnick, S. D. (2024). Spatial memory encoding is associated with the anterior and posterior hippocampus: An FMRI activation likelihood estimation meta-analysis. Hippocampus, 34(11), 575–582. 10.1002/hipo.23632

Suthana, N. A., Ekstrom, A. D., Moshirvaziri, S., Knowlton, B., & Bookheimer, S. Y. (2009). Human Hippocampal CA1 Involvement during Allocentric Encoding of Spatial Information. The Journal of Neuroscience, 29(34), 10512–10519. 10.1523/JNEUROSCI.0621-09.2009

Van Der Ham, I. J. M., & Claessen, M. H. G. (2020). How age relates to spatial navigation performance: Functional and methodological considerations. Ageing Research Reviews, 58, 101020. 10.1016/j.arr.2020.101020

Voldsbekk, I., Bjørnerud, A., Groote, I., Zak, N., Roelfs, D., Maximov, I. I., Geier, O., Due-Tønnessen, P., Bøen, E., Kuiper, Y. S., Løkken, L.-L., Strømstad, M., Blakstvedt, T. Y., Bjorvatn, B., Malt, U. F., Westlye, L. T., Elvsåshagen, T., & Grydeland, H. (2022). Evidence for widespread alterations in cortical microstructure after 32 h of sleep deprivation. Translational Psychiatry, 12(1), 161. 10.1038/s41398-022-01909-x

Wanger, T. J., Janes, A. C., & Frederick, B. B. (2023). Spatial variation of changes in test–retest reliability of functional connectivity after global signal regression: The effect of considering hemodynamic delay. Human Brain Mapping, 44(2), 668–678. 10.1002/hbm.26091

Warren, D. E., Duff, M. C., Tranel, D., & Cohen, N. J. (2011). Observing Degradation of Visual Representations over Short Intervals When Medial Temporal Lobe Is Damaged. Journal of Cognitive Neuroscience, 23(12), 3862–3873. 10.1162/jocn_a_00089

Wechsler, D. (1999). Manual for the Wechsler Abbreviated Intelligence Scale (WASI). The Psychological Corporation.

Wickham, H. (2016). ggplot2: Elegant Graphics for Data Analysis. Springer-Verlag New York. https://ggplot2.tidyverse.org

Xie, W., Chapeton, J. I., Bhasin, S., Zawora, C., Wittig, J. H., Inati, S. K., Zhang, W., & Zaghloul, K. A. (2023). The medial temporal lobe supports the quality of visual short-term memory representation. Nature Human Behaviour, 7(4), 627–641. 10.1038/s41562-023-01529-5

Yarkoni, T., Poldrack, R. A., Nichols, T. E., Van Essen, D. C., & Wager, T. D. (2011). Large-scale automated synthesis of human functional neuroimaging data. Nature Methods, 8(8), 665–670. 10.1038/nmeth.1635

Yonelinas, A. P. (2013). The hippocampus supports high-resolution binding in the service of perception, working memory and long-term memory. Behavioural Brain Research, 254, 34–44. 10.1016/j.bbr.2013.05.030

