## Supplementary Material for "Phase-Specific Hippocampal and Cortical Medial Temporal Lobe Involvement in Allocentric Working Memory"

### Supplemental Material

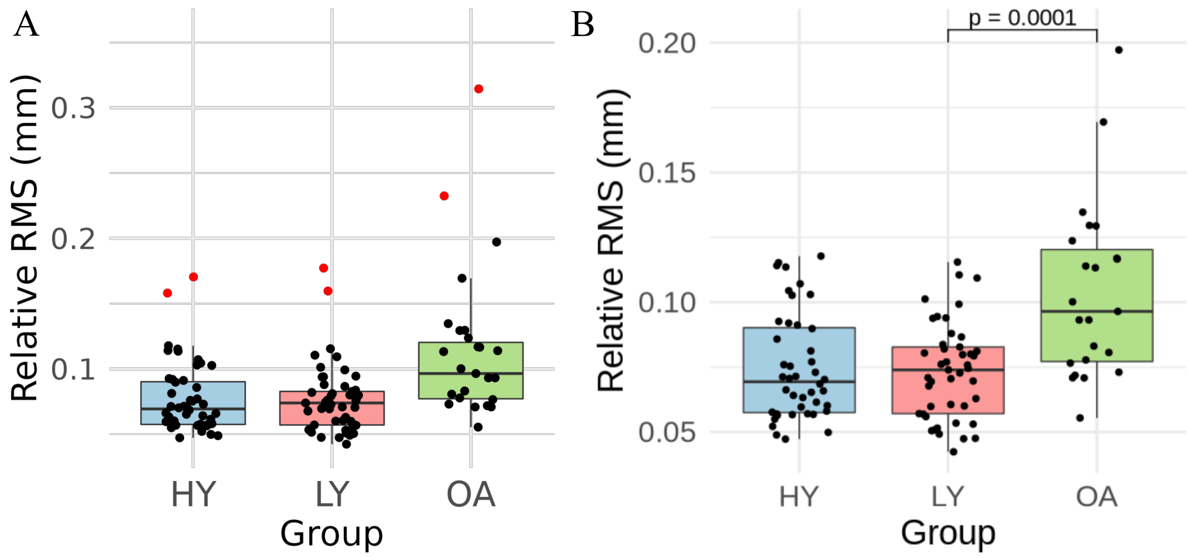


**Figure S1**. Relative root-mean-square (RMS) values by group.
(A) Individual RMS values are shown for higher-performing younger adults (HY), lower-performing younger adults (LY), and older adults (OA). Red points indicate participants who exceeded the exclusion threshold for head motion (1.5 × IQR above the upper quartile), calculated separately within each group.
(B) Despite motion-based exclusions, LY and OA groups still differ significantly in relative RMS values.
